# Tunable control of insect pheromone biosynthesis in *Nicotiana benthamiana*

**DOI:** 10.1101/2022.06.15.496242

**Authors:** Kalyani Kallam, Elena Moreno-Giménez, Ruben Mateos-Fernández, Connor Tansley, Silvia Gianoglio, Diego Orzaez, Nicola J. Patron

## Abstract

Previous work has demonstrated that plants can be used as production platforms for molecules used in health, medicine, and agriculture. Production has been exemplified in both stable transgenic plants and using transient expression strategies. In particular, species of *Nicotiana* have been engineered to produce a range of useful molecules, including insect sex pheromones, which are valued for species-specific control of agricultural pests. To date, most studies have relied on strong constitutive expression of all pathway genes. However, work in microbes has demonstrated that yields can be improved by controlling and balancing gene expression. Synthetic regulatory elements that provide control over the timing and levels of gene expression are therefore useful for maximizing yields from heterologous biosynthetic pathways. In this study, we demonstrate the use of pathway engineering and synthetic genetic elements for controlling the timing and levels of production of Lepidopteran sex pheromones in *Nicotiana benthamiana*. We demonstrate that copper can be used as a low-cost molecule for tightly regulated inducible expression. Further, we show how construct architecture influences relative gene expression and, consequently, product yields in multigene constructs. We compare a number of synthetic orthogonal regulatory elements and demonstrate maximal yields from constructs in which expression is mediated by dCas9-based synthetic transcriptional activators. The approaches demonstrated here provide new insights into the heterologous reconstruction of metabolic pathways in plants.

## Introduction

The reconstruction of biosynthetic pathways in heterologous organisms has become an established route for the production of valuable biomolecules. While recombinant DNA technologies have been in use for decades, recent advances in metabolic engineering and synthetic biology have expanded the breadth and complexity of molecules produced by heterologous biosynthesis (Keating and Young, 2019; Romero-Suarez et al., 2022). Indeed, a major advantage of biological manufacturing is the ability to produce complex molecules, including those for which chemical synthesis has proven difficult or commercially non-viable due to the requirement of multiple stereoselective steps (Cravens et al., 2019). For example, there is growing interest in the biological production of insect sex pheromones, many of which are stereochemically complex, for species-specific control of agricultural pests (Ding et al., 2014; Holkenbrink et al., 2020; Mateos Fernández et al., 2022; Mateos-Fernández et al., 2021; Xia et al., 2021).

Most progress in the heterologous biosynthesis of natural products has been achieved by the engineering of industrially established microbes. However, plant and algal production systems are becoming more widely used (Brodie et al., 2017; Burnett and Burnett, 2020; Stephenson et al., 2020). The use of photosynthetic hosts negates the requirement for sugar feedstocks required by some microorganisms, which, depending on the sources from which they are derived, can raise new issues of sustainability (Dammer et al., 2019; Matthews et al., 2019). Plants can express, fold and post-translationally modify most eukaryotic proteins. They also produce many metabolic precursors and cofactors allowing the facile reconstruction of metabolic pathways often without the need to engineer host genes and pathways (Patron, 2020; Stephenson et al., 2020). For example, precursors of Lepidopteran sex pheromones have been produced in field-grown transgenic *Camelina sativa* (false flax) (Wang et al., 2022)), while biosynthesis of the pheromone molecules has been demonstrated in other plant species (Ding et al., 2014; Mateos-Fernández et al., 2021; Xia et al., 2022).

Tobacco (*Nicotiana tabacum*) and other species in the Nicotiana genus are highly amenable to *Agrobacterium*-mediated transformation and, consequently, have become widely used both as model plants for studying gene function and for biotechnology (Bally et al., 2018; Lein et al., 2008; Molina-Hidalgo et al., 2021). *N. benthamiana*, a non-cultivated species native to Australia, has a comparatively short life cycle and does not accumulate much biomass in field conditions. However, it is particularly amenable to *Agrobacterium*-mediated transient expression (agroinfiltration), which has been exploited for the large-scale production of recombinant proteins, including the production an approved COVID-19 vaccine in Canada (Chen et al., 2013; Hager et al., 2022; Stephenson et al., 2020). In recent years, this method has been shown to be amenable for the reconstruction of many metabolic pathways, including the production of preparative quantities (van Herpen et al., 2010; Molina-Hidalgo et al., 2021; Reed et al., 2017; Stephenson et al., 2020).

Transient production offers many advantages, including a short timeline (less than two weeks) allowing updated construct designs to be rapidly implemented (Chen et al., 2013; Hager et al., 2022; Stephenson et al., 2020). Agroinfiltration also results in the delivery of multiple copies of the construct being delivered to and expressed in all infiltrated cells, enabling high yields. While transplastomic plants leverage the high-copy number of plastid genomes to produce high levels of recombinant proteins, metabolites must be produced in the cellular compartments in which precursors are available. Therefore, transgenic approaches are often limited to nuclear transformation where high copy events often result in gene silencing and low expression. Another advantage of agroinfiltration is that it takes place within contained facilities meaning the lengthy and expensive regulatory processes required for field release of transgenic plants are not required. However, large-scale agroinfiltration has higher energy demands and requires an initial investment in infrastructure. In contrast, transgenic seeds can be easily and cheaply stored and distributed, and transgenic plant lines can be used to produce biomass on an agricultural scale. In particular, *N. tabacum* has been bred for leaf production and accumulates considerable biomass; it has been estimated that field-grown transgenic tobacco are several-fold more cost-effective than cell culture methods for the production of some recombinant proteins (Conley et al., 2011; Schmidt et al., 2019). However, the identification and assessment of high-yielding, stable transgenic lines can be laborious, and, for field growth, regulatory barriers add substantial costs. Further, high-level transgenic expression of some molecules can impact growth and development. For example, we recently showed that growth was compromised in transgenic lines *N. benthamiana* producing the highest yields (per gram fresh weight) of the Lepidopteran sex pheromone components, (Z)-11-hexadecenyl acetate (Z11-16OAc) and (Z)-11-hexadecenol (Z11-16OH) (Mateos-Fernández et al., 2021). Analysis of transcriptional changes revealed stress-like responses, including the downregulation of photosynthesis-related genes (Juteršek et al., 2022). The complex advantages and disadvantages of transient and transgenic approaches make it challenging to determine which strategy will be most cost-effective for large-scale production of a given molecule.

To date, most efforts to express new pathways in plants have used strong constitutive expression of all pathway genes. However, the use of orthogonal synthetic elements reduces the possibility of unpredictable depression resulting from inadvertent interactions with host machinery and has been shown to improve the predictability of engineered circuits (Brophy and Voigt, 2014; Meyer et al., 2019). Further, work in microbial systems has demonstrated that balancing the expression of pathway genes can influence the accumulation of pathway intermediates and precursors and lead to increases in yields (Jones et al., 2015). The availability of characterized regulatory elements and design rules that allow control over expression levels of heterologous pathways, therefore, highly desirable. For example, impacts on growth and development might be overcome by improvements to construct design that allow the timing and levels of expression to be tuned. Further, the ability to tune the relative expression of genes within heterologous pathways might provide the ability to balance metabolic pathways, for example, to control the relative yields of pheromone components, the ratio of which is known to differ between moth species (Zavada et al., 2011).

In this study, we prototype synthetic genetic elements and construct designs for the control of metabolite production in plant systems demonstrating their use in the production of Lepidopteran pheromones. First, we increase the accumulation of (Z)-11-hexadecenyl acetate (Z11-16OAc) using more productive diacylglycerol acetyltransferases. We then demonstrate and compare synthetic regulatory elements assessing their suitability for pheromone production, simultaneously evaluating if transgenic or transient production methods are likely to provide the best net yields. We show that expression systems inducible by copper (Garcia-Perez et al., 2022), a relatively low-cost molecule that is readily taken up by plants and registered for field use (Kumar et al., 2021; Mett et al., 1993; Saijo and Nagasawa, 2014) result in tight control of expression but that highest yields are obtained from a dCas9-based system (dCasEV2.1, (Selma et al., 2019) using transient agroinfiltration. We also demonstrate that construct architectures affect the expression levels of co-assembled synthetic genes in a sequence-dependent manner. We leverage the positional effects on gene expression in multigene constructs to tune the relative levels of the major pheromone components. In addition, we demonstrate that these positional effects are not observed using a synthetic dCasEV2.1 system that allows the use of unique promoter sequences for each enzyme.

## Results

### Comparison of diacylglycerol acetyltransferases

Heterologous production of moth sex pheromones from endogenous 16C fatty acyl CoA is achieved by constitutive expression of Δ11 desaturase, fatty acid reductase and diacylglycerol acetyltransferase (**Figure 1A**). The genes encoding moth acetyltransferases involved in pheromone biosynthesis have not yet been identified and previous attempts at heterologous biosynthesis have employed either an enzyme from the plant, *Euonymus alatus* (EaDAct) (Ding *et al*., 2014; Mateos-Fernández *et al*., 2021), or from the yeast, *Saccharomyces cerevisae* (ScATF1) (Ding *et al*., 2016; Xia *et al*., 2022), with the latter producing more acetate. We first compared EaDAct and ScATF1 with two further diacylglycerol acetyltransferases from *Euonymus fortunii* (EfDAct) *(Tran et al., 2017)* and *Saccharomyces pastorianus* (SpATF1-2) (Yoshimoto *et al*., 1999). To do this, each gene was co-expressed with the coding sequences of enzymes encoding a Δ11 desaturase from *Amyelois transitella* (AtrΔ11) and a fatty acid reductase from *Helicoverpa armigera* (HarFAR) (Figure 1B). We found that both yeast enzymes produced more Z11-16OAc than those from plants (**Figure 1B**).

**Figure 1.**
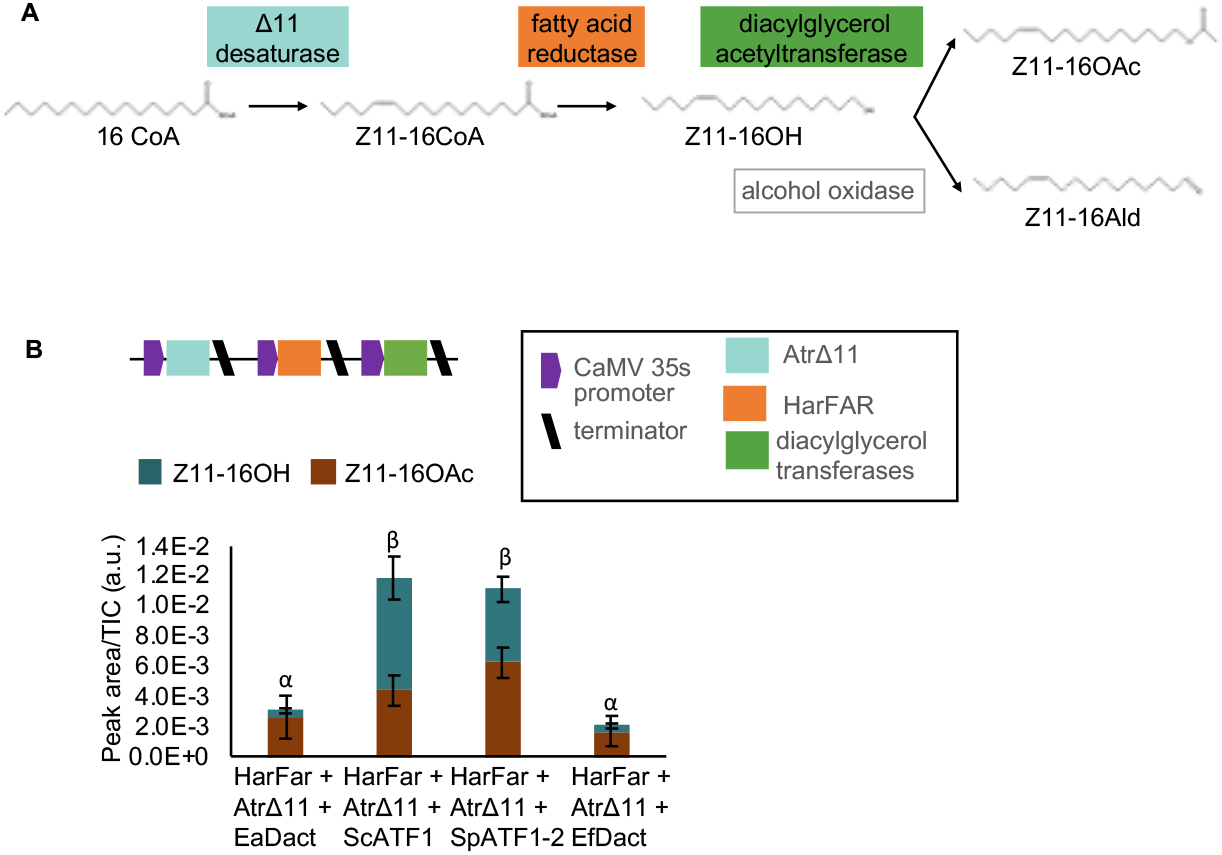
Heterologous production of Lepidopteran sex pheromones. **(A)** Plant production of the two main volatile components in many Lepidopteran sex pheromones *(Z)*-11-hexadecenol (Z11-16OH) and *(Z)*-11-hexadecenyl acetate (Z11-16OAc) from endogenous 16C fatty acyl CoA (Z11-16CoA) was previously achieved by heterologous expression of a Δ11 desaturase, a fatty acid reductase and a diacylglycerol acetyltransferase. The accumulation of *(Z)*-11-hexadecenal (Z11-16:Ald) was also observed, presumably catalyzed by an endogenous alcohol oxidase (Mateos-Fernández *et al*., 2021). **(B)** Differences in the quantities and ratios of Z11-16OAc and Z11-16OH obtained by co-expression of diacylglycerol transferases from *Euonymus alatus (EaDAct), E. fortunii (EfDAct), Saccharomyces cerevisiae* (ScATF1) and *S. pastorianus* (SpATF1-2) with a fatty acid reductase from *Helicoverpa armigera* (HarFAR) and a Δ11 desaturase from *Amyelois transitella* (AtrΔ11). Values shown are the mean and standard error of n=3 biological replicates (independent infiltrations). Means annotated with a common Greek letter (α, β) are not significantly different by a one-way ANOVA with post-hoc Tukey HSD at the 5% level of significance.

### Copper inducible expression of Lepidopteran pheromones

Control over gene expression allows production to be limited to mature plants close to the intended harvest time, limiting effects on plant growth. Tightly controlled inducible regulatory systems are useful tools. However, to reach the scales required for cost-effective production, any agents used to induce expression must be low-cost and, ideally, usable in open-field systems. In previous work, we showed that synthetic transcriptional activators comprised of translational fusions of the yeast protein, CUP2, which binds to cognate DNA sequences in the presence of copper (Buchman et al., 1989), and the yeast transcriptional activator, Gal4, (Ma and Ptashne, 1987) resulted in strong, copper-inducible activation of minimal synthetic promoters containing CUP2 binding sites (CBSs) (Garcia-Perez et al., 2022). We therefore tested copper-inducible accumulation of pheromone components, reasoning that copper sulfate is low-cost and already used in agriculture. To do this, we assembled the coding sequences of AtrΔ11, HarFAR and SpATF1-2 with a minimal 35s promoter preceded by four copies of the CBS (**Figure 2A**). These three synthetic genes were then co-assembled with a synthetic gene in which the *A. tumefaciens* nopaline synthase promoter (AtuNos) was fused to CUP2:GAL4 for moderate constitutive expression (**Figure 2A**). The resulting multigene construct (construct 678) was agroinfiltrated into *N. benthamiana* leaves in a 1:1 ratio with an *Agrobacterium* strain carrying a plasmid encoding the P19 suppressor of silencing (Garabagi et al., 2012). Three days post-infiltration, leaves were sprayed with either water or 2.5 mM copper sulfate (CuSO_4_), previously identified as the optimal concentration (Garcia-Perez et al., 2022). The total volatile organic compound (VOC) composition of all samples was analyzed five days post-infiltration by gas chromatography/mass spectrometry (GC/MS). GC peaks corresponding to the pheromone compounds Z11-16OH and Z11-16OAc were detected in samples treated with CuSO_4_, but not in untreated samples or in control samples infiltrated with P19 alone (**Figure 2B**). The best yield obtained was 12.4 μg Z11-16OH g^-1^ fresh weight (FW) and 4.5 μg Z11-16OAc g^-1^ FW.

**Figure 2.**
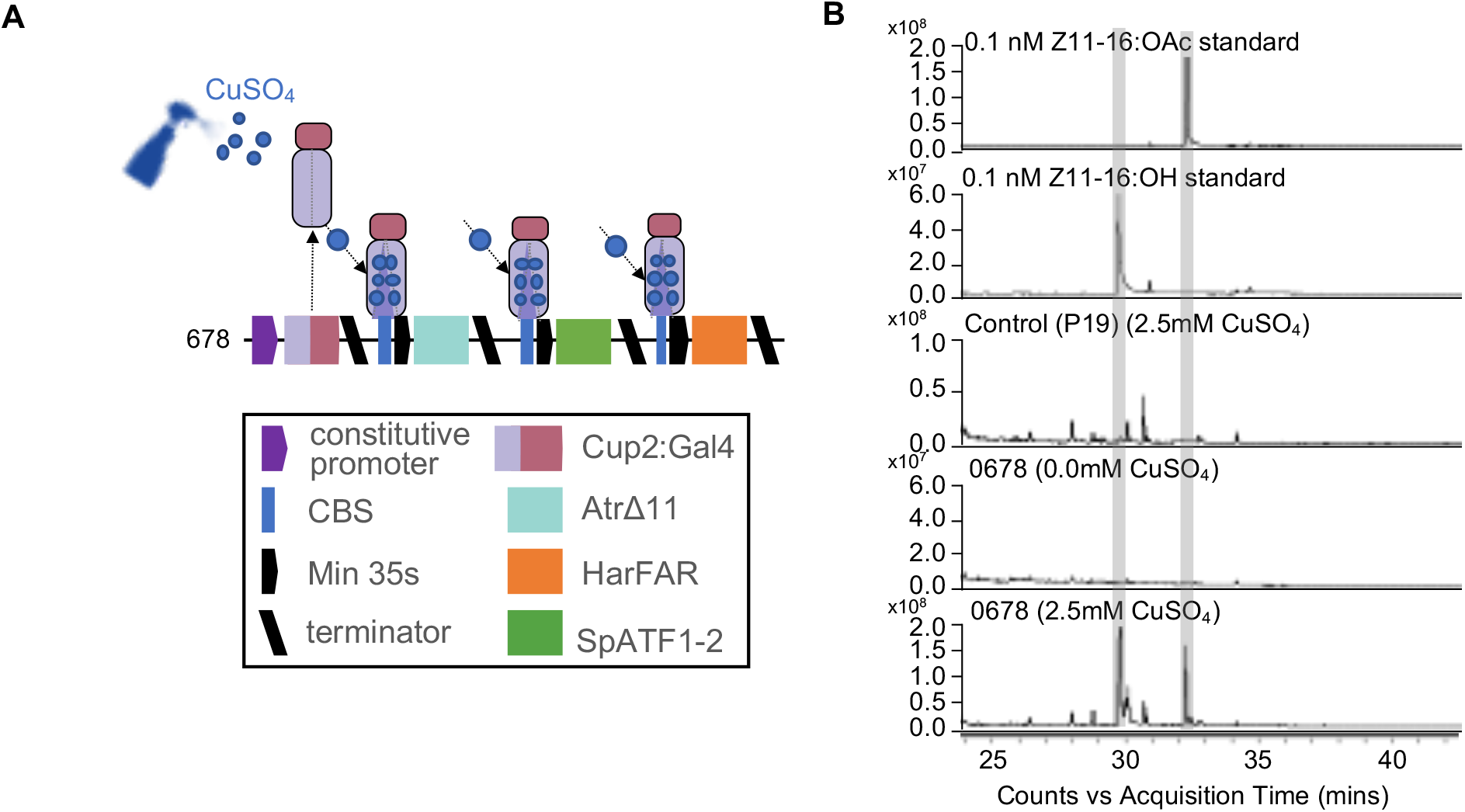
Copper inducible expression of Lepidopteran pheromones. (**A)** Schematic of a plant expression construct containing synthetic genes encoding the copper-responsive transcription factor CUP2 in translational fusion with the Gal4 activation domain and the coding sequences of *AtrΔ11, HarFAR* and *SpATF1-2* under control of a minimal 35s promoter preceded by four copies of the CUP2 binding site (CBS). (**B)** Total ion chromatogram showing the accumulation of Z11-16:OH and Z11-16:OAc in leaves of *N. benthamiana* co-infiltrated with *Agrobacterium* strains containing the expression construct (678) and a construct expressing the P19 suppressor of silencing only after application of 2.5 mM copper sulfate (CuSO_4_).

### Construct architecture influences expression and product yield

It has long been known that the repetition of some genetic elements within constructs as well as the insertion of T-DNA as tandem repeats can trigger gene silencing (Stam *et al*., 1997; Vaucheret *et al*., 1998). This presents a challenge for designing synthetic circuits in which coordinated expression of multiple genes in response to a single signal is desirable. For transient expression, it is possible to avoid co-assembly onto a single T-DNA by the co-delivery of multiple strains of *A. tumefaciens*. However, it is unknown what proportion of cells receive all strains and if this affects maximum yields. Further, when producing stable transgenic lines, it is desirable that pathway genes are coassembled to enable integration into a single genomic locus, preventing segregation in the progeny.

To determine if and how expression levels of synthetic genes are affected by co-assembly, and to investigate if adjustments to construct architecture would affect yield and product ratios, we first compared expression in single and multigene assemblies. Initially, relative expression levels from two luciferase reporters driven by CaMV35s promoters in single and multi-gene configurations were compared (**Figure 3A**). In multigene configurations, transcription units were assembled on the same strand; constructs in which the two genes were assembled on opposing strands were unstable. We observed that coinfiltration of separate constructs expectedly resulted in equal quantities of each reporter, however, the relative expression within multi-gene constructs was affected by the position of the gene in the assembly, with relatively more expression from the first gene (**Figure 3A**). To investigate if the same effect is seen with copper-inducible regulatory promoters, we performed equivalent assays with two versions of copper-inducible synthetic promoters.

**Figure 3.**
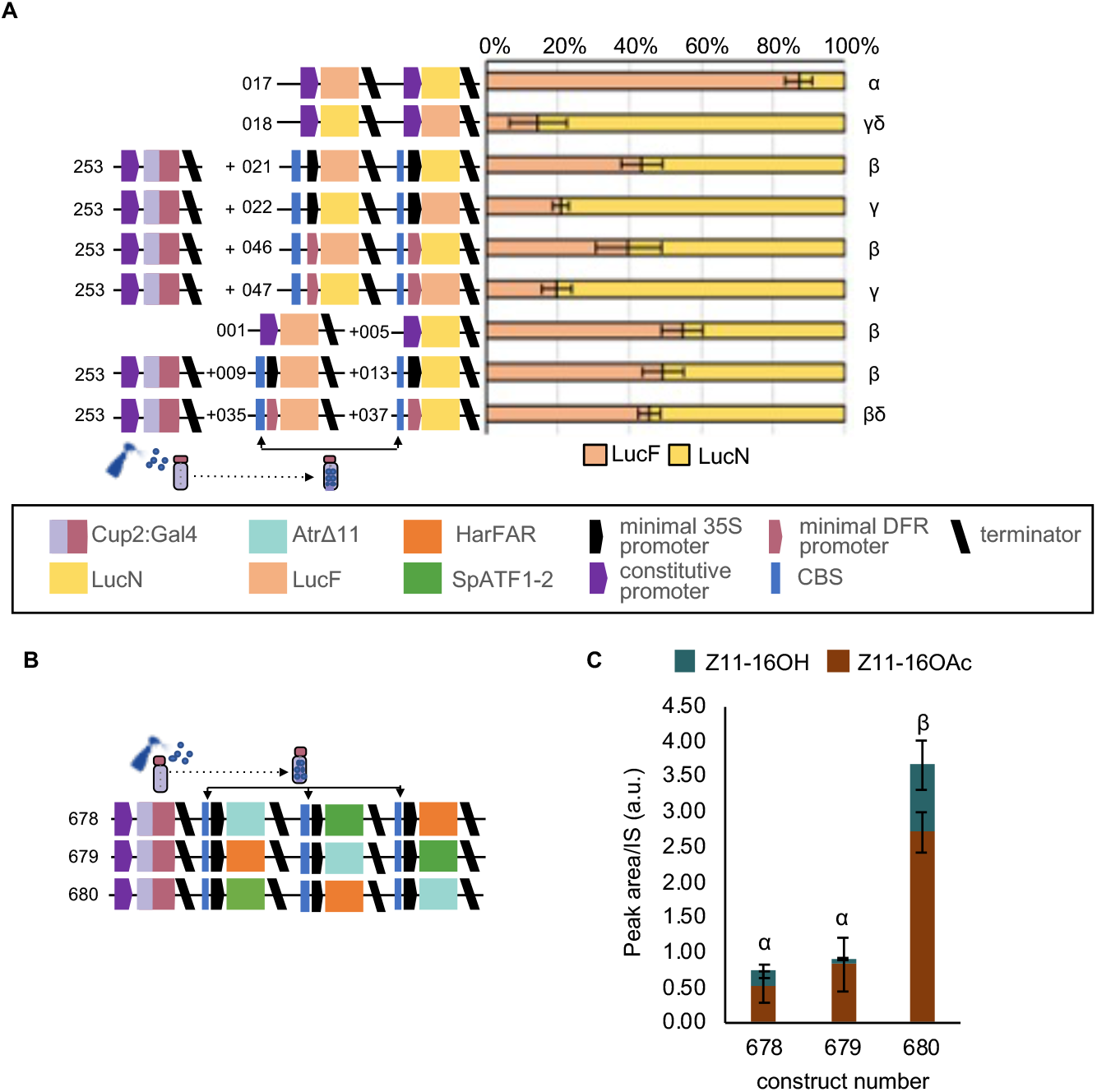
Construct architecture influences expression and product yield. (**A)** The level of expression of firefly luciferase (LucF) and nanoluciferase (LucN) in a multigene construct is dependent on the position in which the gene is assembled. Values shown are the mean and standard error of n=6 biological replicates (independent infiltrations) and differences were analyzed using a Kruskal-Wallis test followed by pairwise Wilcoxon rank sum test with Benjamini-Hochberg correction. Bars annotated with a common Greek letter (α, β, γ, δ) are not significantly different. (**B**) Schematics of plant expression constructs containing synthetic genes for copper-inducible expression of lepidopteran sex pheromones. (**D**) The relative positions of pathway genes influenced the overall yield and the relative ratios of pheromone products. Values shown are the mean and standard error of n=3 biological replicates (independent infiltrations). Means annotated with common Greek letters (α, β) are not significantly different by a one-way ANOVA followed by Post-hoc Tukey test at the 5% level of significance.

The first version used a minimal 35s promoter preceded by four copies of the CBS. In the second version, the minimal 35s promoter was replaced with a minimal promoter of the *Solanum lycopersicum* NADPH-dependent dihydroflavonol reductase (DFR) (Garcia-Perez *et al*., 2022). In both cases, luminescence was greater in leaves treated with CuSO_4_, however, while background expression in the absence of CuSO_4_ was somewhat lower with the minimal DFR, higher expression was obtained with the minimal 35s promoter (Supplementary Figure 1). Although less pronounced than with the CaMV35s promoters, expression levels obtained from copper inducible genes were also affected by co-assembly into multigene constructs (**Figure 3A**). This data demonstrates that relative expression levels measured for individual synthetic genes are not always maintained in multigene constructs, and that the effects are sequence dependent.

From these results, we reasoned that altering the relative position of genes in constructs encoding the pheromone biosynthesis pathway would likely affect relative expression and thus alter the accumulation of total and relative quantities of Z11-16OH and Z11-16OAc. To investigate this, we assembled and compared three copper-inducible constructs within which we varied the relative positions of each gene (**Figure 3B**). Consistent with our observation of reporter genes, we observed variations in both the overall yields and the relative ratios of Z11-16OH and Z11-16OAc components (**Figure 3C**). The construct configuration with *AtrΔ11* in the last position (construct 680) improved yields threefold.

### Copper inducible CRISPR-mediated control of gene expression

The CUP2:GAL4 transcriptional activation system was previously used to control expression of a CRISPR-based programmable activator, enabling tightly-regulated control of the expression of both synthetic and endogenous genes (Garcia-Perez et al., 2022). In recent years several orthogonal synthetic activators have been demonstrated in plants but have not been directly compared. To determine which synthetic promoters might provide the best levels of activation and background expression levels when combined with copper-inducibility, we compared three previously reported synthetic promoters activated by (i) a transcription activator-like effector (TALE) (Cai et al., 2021), (ii) a Gal4:ΦC3 fusion protein (Bernabé-Orts et al., 2020; Cai et al., 2021), and (iii) the dCasEV2.1, which consists of dCas9 fused to the EDLL transcriptional activation domain (Cas9:EDLL), the MS2 phage coat protein fused to a synthetic VPR transcriptional activation domain (MS2:VPR), and a guide RNA (gRNA) that guides the complex to a recognition sequence in the promoter (Selma et al., 2019). Expression of the all proteincoding elements was controlled by a copper inducible promoter, except for the expression of the gRNA, which is controlled by the RNA polymerase III dependent Arabidopsis promoter, U6-26, previously demonstrated to function in *N. benthamiana* (Castel et al., 2019; Garcia-Perez et al., 2022). All systems were functional with expression increasing with the application of CuSO_4_ and with the number of binding sites in the synthetic promoter (**Figure 4 A-C**). However, although the maximal expression levels obtained from the TALE and dCasEV2.1 system were similar (**Figure 4B and C**), the expression levels from the TALE system in the absence of copper were considerable and only the dCasEV2.1 system retained low levels of background expression in the absence of copper (**Figure 4C**).

**Figure 4.**
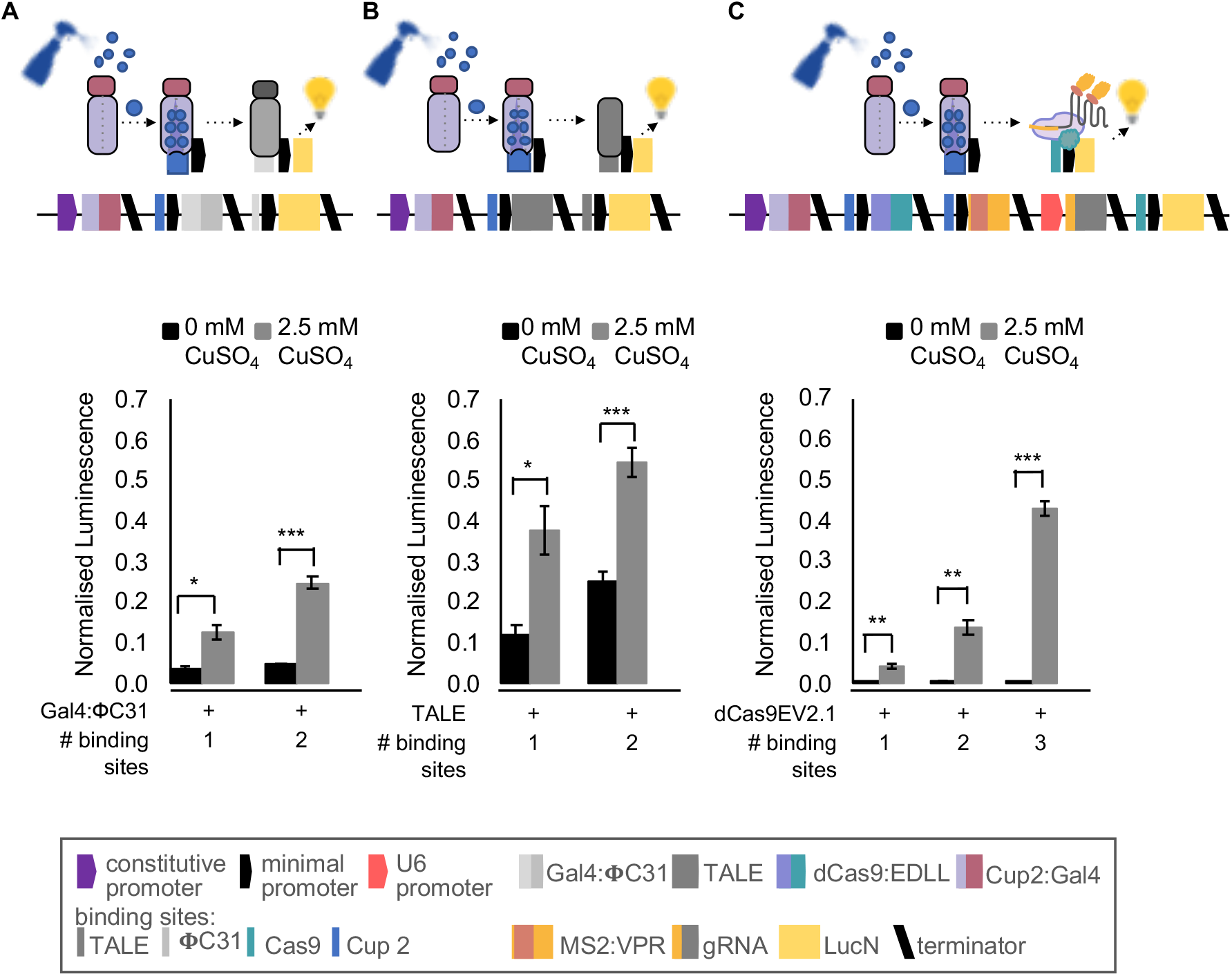
Comparison of synthetic transcriptional activators. Normalized luminescence from reporter constructs activated by copper-inducible **(A)** GAL4:ΦC31 (**B**) activator-like effector (TALE) and (C) dCasEV2.1 synthetic transcriptional activators. In all systems, expression levels increase with copper and with the number of transcriptional activator binding sites in the promoter. Copper inducible expression of dCasEV2.1 maintains tight control (low background) of gene expression. Values shown are the mean and standard error of n=3 biological replicates (independent infiltrations). P-values were calculated using Welch two sample *t-test: *P≤ 0.05, **P≤ 0.01, ***P ≤ 0.001*.

To investigate if this copper-sensing dCasEV2.1 system would enable control over pheromone biosynthesis, we assembled the coding sequence of each pathway enzyme with a synthetic promoter activated by the dCasEV2.1 system. To reduce the amount of sequence repeated within each transcriptional unit, and therefore minimize the potential for any gene silencing mediated positional effects, pathway genes were each assembled with synthetic promoters that had minimal sequence similarity: Each promoter consisted of a unique sequence with the only repeated sequences being the recognition sites for the gRNA and the minimal DFR core region.

This design minimized sequence repetition within the multigene assembly while maintaining activation to a single transcriptional activator. As previously, three assemblies were produced, altering the relative position of each pathway gene (**Figure 5A**). These constructs were coinfiltrated with the copper-sensing dCasEV2.1 module (GB4070). In contrast to the direct copper activation, in which yields were affected by construct architecture (**Figure 3C**), all constructs produced similar ratios of pheromone components, with slightly more Z11-16OH than Z11-16OAc (**Figure 5B**), indicating that these regulatory elements were less affected by co-assembly. Yields obtained using the copper-sensing dCasEV2.1 reached 32.7 μg Z11-16OH g^-1^ FW and 25 μg Z11-16OAc g^-1^ FW. We also repeated the entire experiment replacing SpATF1.2 with ScATF1, with similar results (Supplementary Figure S2).

**Figure 5.**
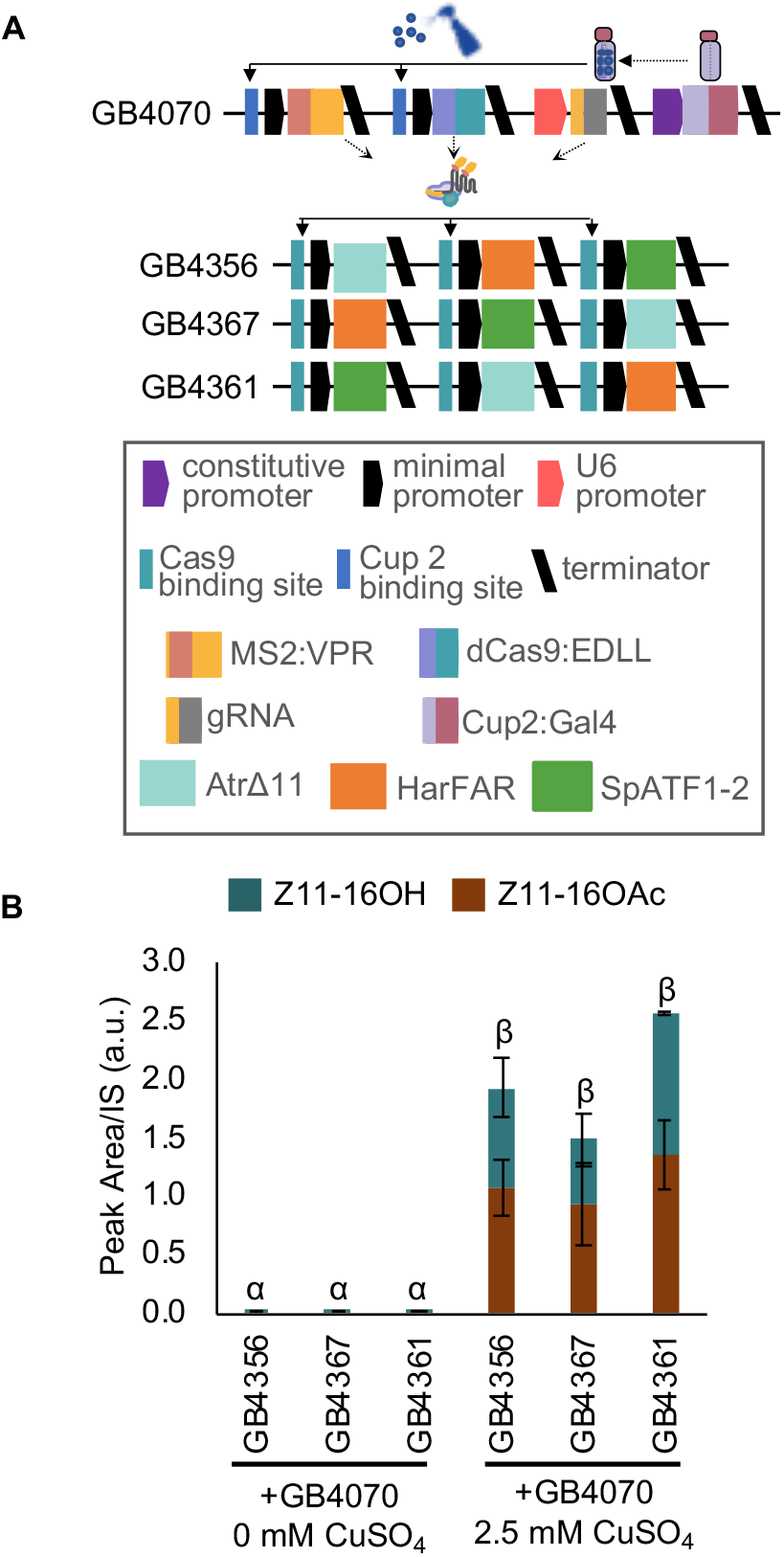
Copper inducible, CRISPR/Cas9-mediated control of pheromone biosynthesis. (**A)** Schematic of plant expression constructs containing elements for copper inducible expression of the dCasEV2.1 transcriptional activator (above) and multigene constructs containing coding sequences for AtrΔ11, HarFAR and SpATF1-2, each assembled with a unique promoter with cognate target sequences for the gRNA (**B)** Application of CuSO_4_ results in dCasEV2.1 mediated production of the pheromone components (Z11-16OH and Z11-16OAc). Values shown are the mean and standard error of n=3 biological replicates (independent infiltrations). Means followed by a common Greek letter (α, β) are not significantly different (one-way ANOVA with post-hoc Tukey HSD at the 5% level of significance).

### Copper-inducible expression in stable transgenics

The above transient experiments indicate that copper inducible synthetic elements enable tight control of heterologous pathway genes. This would enable expression to be induced after the accumulation of biomass, potentially limiting effects on plant growth. However, expression levels and pheromone yields obtained from copper-inducible promoters were observed to be reduced as compared to those achieved from constitutive promoters. As the copy number and, therefore, yield is also expected to be reduced in stable transgenics, it is important to quantify potential expression levels in such lines. To investigate expression levels from the copper sensing dCasEV2.1 system in stable transgenics, we produced plants expressing three the regulatory components: dCas9:EDLL and MS2:VPR under the control of copper-inducible promoters and the CUP2:GAL4 transcriptional activator under the control of the constitutive nopaline synthase (nos) promoter (Figure 6A). The resulting plant lines provide a modular, reusable resource that could be crossed with lines expressing synthetic pathways driven by orthogonal promoters with binding sites for one or more co-expressed single guide RNAs (sgRNA). To identify high-performing lines, we infiltrated ten independent T_0_ plants with constructs encoding firefly luciferase (LucF) under the control of the previously tested synthetic promoter with three recognition sites for the gRNA and the gRNA, together with a constitutively expressed Renilla luciferase (LucR) calibrator gene. Three leaves of each plant were infiltrated and 0.0 mM CuSO_4_ or 2.5 mM CuSO_4_ were applied to each side of the midrib. Protein was extracted and dual luciferase assays were used to quantify expression. Expression was compared to non-transgenic lines in which all components were transiently expressed (Figure 6B). One line, CBS:dCas4, was identified in which 2.5 mM CuSO_4_ resulted in a significant increase in expression. Expression from stable transgenics was considerably less (~85 fold) than from plants in which all constructs were transiently expressed, presumably due to the reduced availability of dCasEV2.1 components (Figure 6B). To confirm this, T1 seed from three lines were collected and RNA was extracted from plants treated with 0.0 mM CuSO_4_ or 2.5 mM CuSO_4_. The expression levels of dCas9:EDLL and MS2:VPR were quantified by qRT-PCR, finding that mRNA levels correlated with luminescence (Supplementary Figure S3). These data indicate that, while the copper-sensing dCasEV2.1 system is functional when integrated into the plant genome, that stable transgenic lines are unlikely to produce useful levels of pheromones.

**Figure 6.**
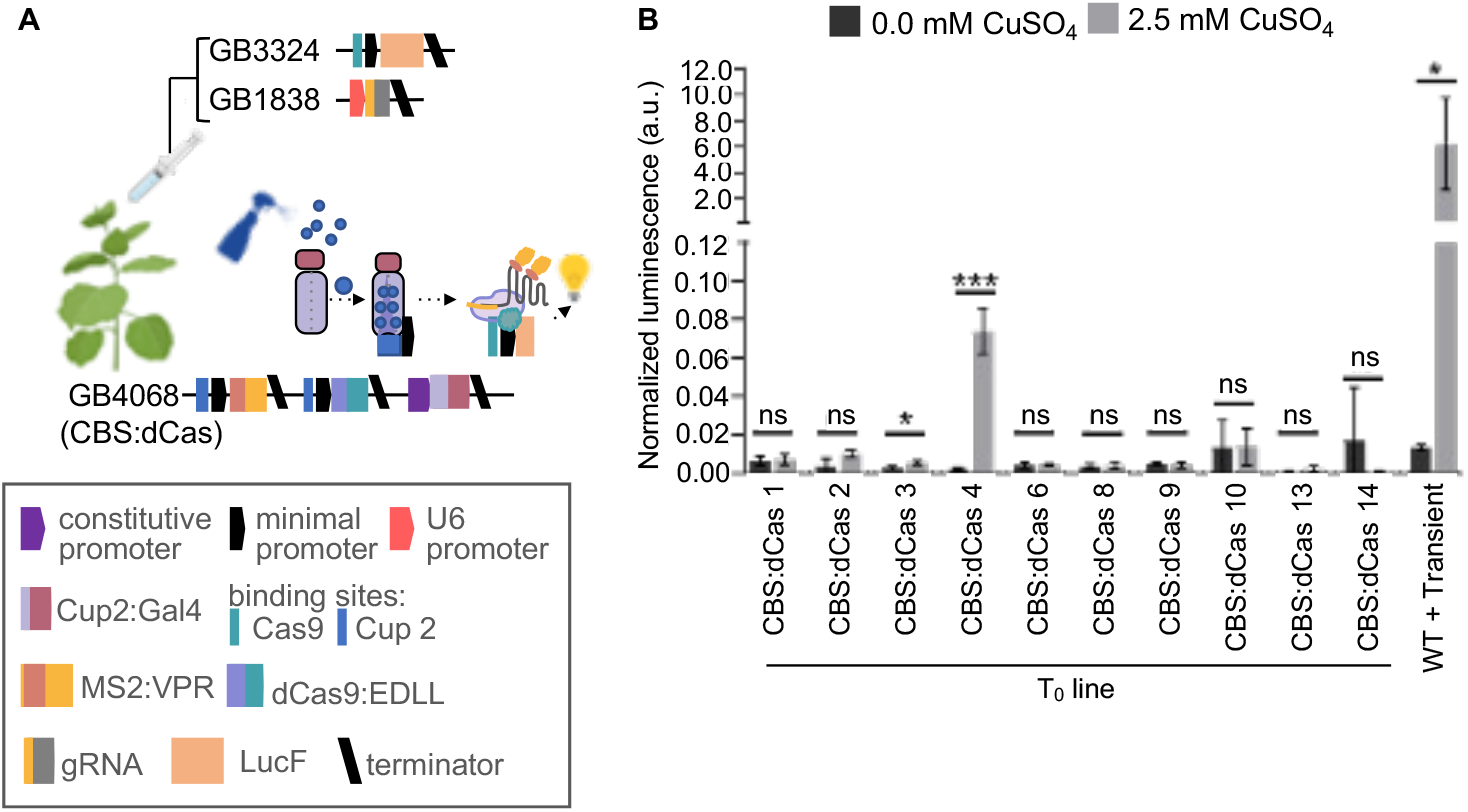
Functionality of the copper sensing dCasEV2.1 module in stable *Nicotiana benthamiana* transgenics. (**A**) Transgenic plants expressing the CBS:dCasEV2.1 (construct GB4068) were agroinfiltrated with a luciferase reporter module (construct GB3324) and sgRNA module (construct 1838). (**B**) Normalized expression levels of luciferase in T_0_ CBS:dCas transgenic plants after copper induction. Values are the mean and standard deviation of n=3 independent infiltrations. P-values were calculated using Student’s *t-test; * P ≤ 0.05, *** P ≤ 0.001;* ns= not significant. The figure includes images from Biorender (biorender.com).

### CRISPR-mediated control of pheromone biosynthesis

The experiments with dCasEV2.1 demonstrate that this system enables the construction of multigene pathways with minimal repetition of regulatory sequences sequence repetition while maintaining activation to a single transcriptional activator (**Figure 5**). We therefore assessed the yields obtained from constitutively expressed dCasEV2.1 activated pheromone pathways using transient agroinfiltration. We compared two construct configurations in which the gRNA element was either coassembled with the dCasEV2.1 elements (constructs GB2513 + GB3898, non-guided) or the pathway genes (constructs GB2085 + GB3897, guided) (**Figure 7A**). Both configurations were functional, and yields were comparable to those obtained from the CaMV35s promoter (construct GB4407) (**Figure 7B**). The best yields were obtained from construct GB3898 + GB2513 (384.4 μg Z11-16OH g^-1^ FW and 175.8 μg Z11-16OAc g^-1^ FW). Data obtained from copper-inducible transgenic lines suggested the abundance of transcriptional activators was critical. To investigate this further, we created transgenic lines encoding the guided and non-guided dCasEV2.1-activatable pathway (constructs GB3897 and GB3998) (**Figure 7C**). Twenty-five T_0_ plants (7 guided and 18 nonguided) were infiltrated with dCasEV2.1 regulatory elements and pheromone content was assessed (Supplementary Figure S4). Four pheromone-accumulating lines were selected and pheromone accumulation following infiltration of dCasEV2.1 regulatory elements was quantified in T_1_ progeny (**Figure 7D**). The maximum yields obtained were 12.5 μg Z11-16OH g^-1^ FW and 2.8 μg Z11-16OAc g^-1^ FW from line NGP 38. Considering the reduced copy number of the pathway genes compared to transient infiltration, these yields are reasonable and confirm that availability of transcriptional activators is critical. Transgenic production of metabolites may, therefore, be feasible, but will require a system that enables inducible, high-level expression of the dCasEV2.1 elements.

**Figure 7.**
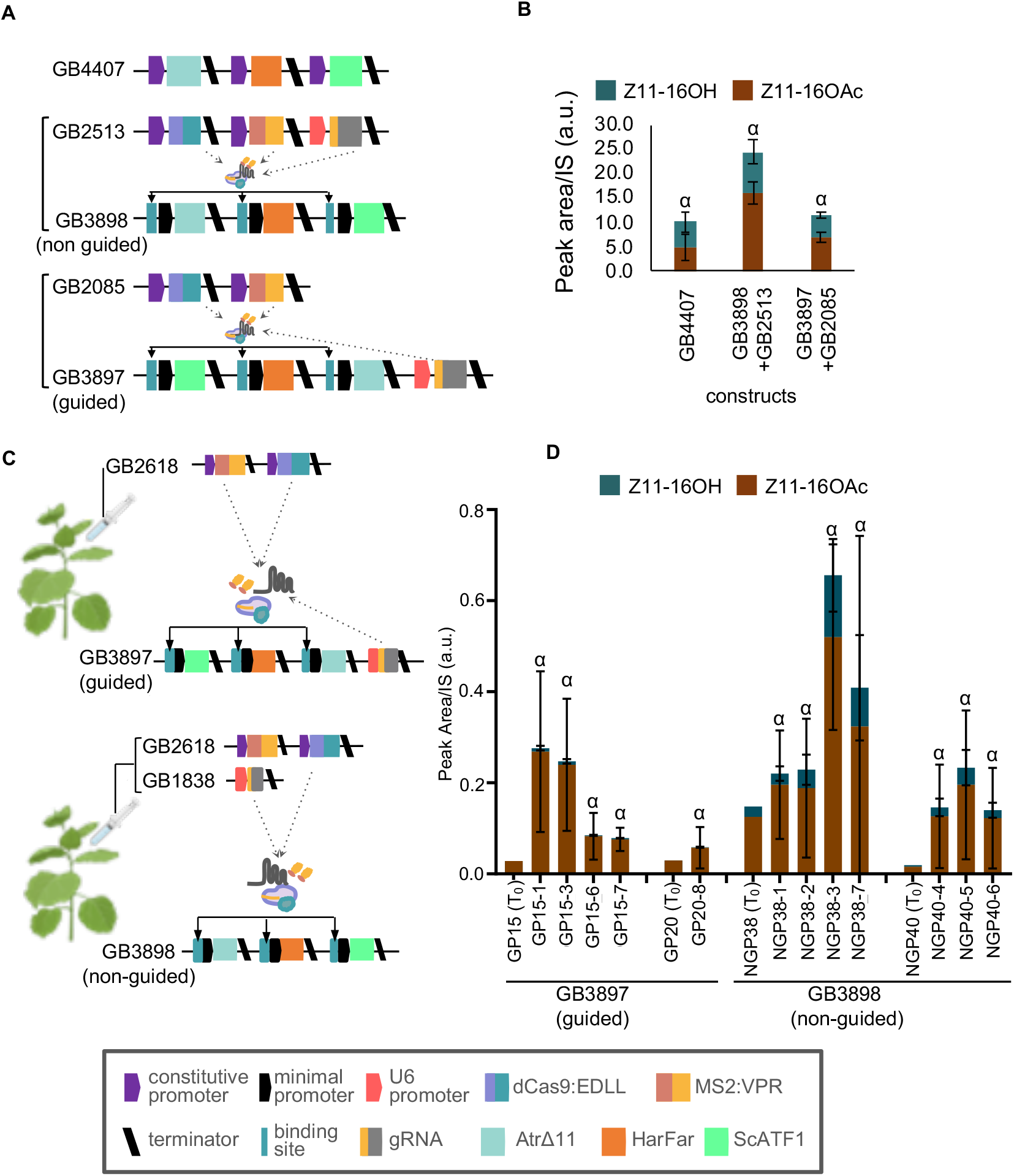
CRISPR/Cas9-mediated control of pheromone biosynthesis. (**A**) Schematic of constructs for constitutive or dCasEV2.1 activated expression of AtrΔ11, HarFAR and SpATF1-2 (**B**) Yields of pheromone components (Z11-16OH and Z11-16OAc) obtained following transient agroinfiltration. (**C**) Schematics of constructs encoding the guided pathway (sgRNA integrated) or non-guided pathway (sgRNA infiltrated) used to produce transgenic lines and transiently expressed dCasEV2.1 activating elements (**D**) Pheromone levels obtained from T_1_ progeny of four independent T_0_ lines encoding either the guided (GB2618) or non-guided (GB3898) pathway infiltrated with constructs expressing regulatory elements. Values represent the mean and standard deviation of n=3 biological replicates (independent infiltrations). Values annotated with a common Greek letter (α) are not significantly different (one-way ANOVA with post-hoc Tukey HSD at the 5% level of significance). This figure includes images from Biorender (biorender.com).

## Discussion

Plants, particularly species of *Nicotiana* are emerging as useful platforms for heterologous production of a range of small molecules for health, industry and agriculture (Brückner and Tissier, 2013; Dudley et al., 2022; van Herpen et al., 2010; Mateos-Fernández et al., 2021; Mikkelsen et al., 2010; Reed et al., 2017; Schultz et al., 2019; Stephenson et al., 2020). Metabolic engineering in microbial systems has demonstrated that optimization of expression constructs to balance pathways and engineering host metabolism can strongly influence yields, (Jensen and Keasling, 2015; Jones et al., 2015). In particular, the use of orthogonal synthetic elements has been shown to limit unpredictable expression patterns and to improve the predictability of engineered circuits (Brophy and Voigt, 2014; Meyer et al., 2019). Such elements also provide the ability to control the timing and levels of expression to limit impacts on growth and development, as well as the ability to tune the relative expression of genes within heterologous pathways to enable pathway balancing. However, to date, heterologous pathway reconstruction in plants has largely been limited to constitutive overexpression.

In this study we used a number of synthetic genetic regulatory elements to control the production of moth sex pheromones. The use of chemical formulations to control insect pests in food crops has a long history. However, the increasing use of synthetic pesticides in the twentieth century led to concerns about the deterioration of biodiversity in agricultural landscapes, as well as risks to farmworkers and consumers (Köhler and Triebskorn, 2013). One alternative is expanding the use of insect sex pheromones, volatile molecules typically produced by females to attract a mate, of which even minute quantities from alternative sources can disrupt breeding and behavior (Mateos Fernández et al., 2022). However, while the pheromones of some species can be cheaply manufactured by synthetic chemistry, the pheromones of many insect species have complex structures requiring stereoselective steps making them difficult and expensive to produce (Petkevicius et al., 2020). As proof-of-concept for the biological synthesis of pheromone components, several studies have successfully produced Lepidopteran sex pheromones in heterologous systems including yeast (Hagström et al., 2013; Holkenbrink et al., 2020; Konrad et al., 2017; Petkevicius et al., 2020) and *N. benthamiana* (Ding et al., 2014; Mateos-Fernández et al., 2021; Xia et al., 2022, 2020). The genes encoding moth acetyltransferases involved in pheromone biosynthesis have not yet been identified and previous studies have employed either a plant enzyme, EaDAcT (Ding et al., 2014; Mateos-Fernández et al., 2021) or the yeast enzyme ScATF1 (Ding et al., 2016; Xia et al., 2022). Here we compared the activities of these enzymes together with an additional enzyme, SpATf1-2 (**Figure 1**), finding that the yeast enzymes produced more acetate. We then used these enzymes for pathway reconstruction with a number of different regulatory elements to enable transient and transgenic expression in *N. benthamiana*.

Both transient and transgenic approaches have been demonstrated for expressing heterologous molecules. Both strategies are, in principle, capable of being scaled for large-scale production. Large-scale transient expression requires more costly infrastructure than field-growth of transgenic lines and, being limited to young plants, cannot achieve high-biomass at low production cost. However, agroinfiltration has a short timeframe, high copy-number (and therefore yield per gram of biomass). In addition, there is no requirement to identify and characterize specific high-yielding lines, or for the expensive regulatory approval required for field growth. The economics of scaling-up production are likely to be different for different molecules, however, there is merit in comparing production methods. In previous experiments, we observed that the growth of transgenic lines of *N. benthamiana* producing the highest yields (per gram fresh weight) of the moth pheromones was negatively affected and photosynthesis-related genes were downregulated (Juteršek et al., 2022; Mateos-Fernández et al., 2021). Inducible gene expression systems are essential tools commonly employed to switch metabolism from growth to production, enabling biomass to accumulate before energy is redirected into the biosynthesis of desired products. Many inducible systems for controlling gene expression in plants use expensive molecules such as estradiol or dexamethasone, or require the application of stresses such as heat or wounding, which cannot easily be applied to large numbers of plants and may affect plant fitness (Corrado and Karali, 2009). However, expression systems inducible by copper, a relatively low-cost molecule that is readily taken up by plants and registered for field use, have also been demonstrated (Kumar et al., 2021; Mett et al., 1993; Saijo and Nagasawa, 2014). In previous work, we optimized a copper-inducible system demonstrating that this enabled high levels of expression of reporter and endogenous genes in *N. benthamiana* and, in the absence of copper, very low background expression (Garcia-Perez et al., 2022). The range of concentrations at which copper is active as a signaling molecule is much lower than those employed for antifungal applications in field conditions, therefore the employment of copper sulfate as a trigger for recombinant gene expression could be compatible with the current reduction trend in copper-based antifungal formulations. Here, we demonstrate that this system is suitable for controlling the expression of biosynthetic pathways (**Figure 2**). As copper sulfate is approved for agricultural use, it provides a possible tool for large-scale bioproduction systems.

The development of modular cloning systems such as Golden Braid (2013; Vazquez-Vilar et al., 2017), used in this study, has eased the assembly of constructs. It is now relatively easy to build variants of genetic constructs that enable the properties of individual components such as promoters and untranslated regions to be compared (Bernabé-Orts et al., 2020; Cai et al., 2021). These cloning systems have also facilitated the design and assembly of multigene constructs (Pollak et al., 2019; Sarrion-Perdigones et al., 2011; Weber et al., 2011). However, few studies have investigated how co-assembly affects the performance of synthetic genes. While it has long been known that gene-silencing can reduce expression from transgenes and that some regulatory elements and construct architectures (e.g. the inclusion of inverted repeats) are more susceptible (Stam et al., 1997; Vaucheret et al., 1998), the effects of co-assembly on individual genes within multigene plant constructs have not been quantified. Using ratiometric reporter assays, we found that the position in which genes are located within multigene constructs differentially affects their expression (**Figure 3**). In other systems, the dominance of the upstream genes has been proposed to be caused by positive supercoiling accumulates downstream of constitutively active genes, altering RNA polymerase binding and initiation (Johnstone and Galloway, 2022). In our work, we found the extent to which genes were affected by co-assembly to be sequence-dependent (**Figure 3**). Extensive studies will be required to investigate how genes in large and complex constructs behave and to determine the best construct architectures for multigene constructs. This will be especially important as more information emerges about the impact of different regulatory elements, including the effects of untranslated sequences and terminators on expression and post-transcriptional silencing (F de Felippes et al., 2020; Wang et al., 2020). It may also be possible to achieve more equal levels of expression by testing the efficacy of insulator sequences that have been used to reduce the effects of genomic locations on transgene expression (Pérez-González and Caro, 2019). In this study, rather than attempting to avoid the unequal expression from coassembled copper-inducible genes, we investigated whether the different levels of expression could be used to alter the product profile obtained from our biosynthetic pathway. We found that changing the relative position of each pathway gene within the construct altered both overall yield and the relative quantities of the major pheromone components (**Figure 3B**).

We also coupled copper-inducible to a CRISPR-based programmable activator, dCas9EV2.1, previously shown to enable tightly-regulated upregulation of endogenous genes (Garcia-Perez et al., 2022). Compared to TALE and PhiC3-based synthetic regulatory elements, dCas9EV2.1 had low to undetectable background expression in the absence of copper (**Figure 4**). Further, the gRNA binding sites could be positioned within different unique promoter sequences to avoid repeating sequence elements within multigene constructs. The yield from these assemblies was comparable to direct use of copper-inducible promoters but were not affected by combinatorial rearrangements (**Figure 5**), confirming that positional effects are sequence-dependent and identifying an alternative approach for avoiding positional effects in multigene constructs.

In previous studies we observed that constitutive expression impacts biomass (Juteršek et al., 2022; Mateos-Fernández et al., 2021). Although yields obtained using copper inducible promoters in transient infiltration were lower (12.4 μg Z11-16OH g^-1^ FW and 4.5 μg Z11-16OAc g^-1^ FW), than those that we obtained in previous studies using CaMV35s promoters (116.6 μg Z11-16OH g^-1^ FW and 110.1 μg Z11-16OAc g^-1^ FW), a lower yield per unit of biomass might be compensated for by high biomass production. We therefore assessed how expression levels from copper inducible promoters in stable transgenics compared to transient agroinfiltration. We found that transgenic lines expressing the copper inducible elements had up to 85-fold reduction in expression (**Figure 6**). We do not consider these expression levels to be viable for pheromone production and conclude that alternative field-compatible, inducible expression systems must be tested or developed. Pheromone yields from alternative species in which production is limited to specific organs might also be tested. For example, precursors of Lepidopteran sex pheromones have recently been produced in the seeds of field-grown transgenic Camelina sativa (false flax) (Wang et al., 2022).

Following a comparison of three orthogonal regulatory systems (**Figure 4**), we also tested pheromone production using dCas9EV2.1 regulatory elements (**Figure 7**). This resulted in the highest yields (384.4 μg Z11-16OH g^-1^ FW and 175.8 μg Z11-16OAc g^-1^ FW). As noted above, this system had the additional advantage of using unique promoter sequences while maintaining activation to a single transcriptional activator and which negated positional effects (**Figure 5**). Finally, we tested dCas9EV2.1 activation of stable transgenes to investigate if maintaining an abundance of transcriptional activators could maintain high yields. The maximum yields obtained from T1 lines were approximately reduced ~20-fold, suggesting that transgenic production may be viable if coupled with a system to enable high level, inducible expression of dCas9EV2.1 elements.

The potential of plants as living biosensors of pheromones has previously been discussed (Mateos Fernández et al., 2022) and trichome-specific promoters have recently shown to increase the release of pheromones from leaves (Xia et al., 2022). However, pheromone components can be extracted from plant biomass for use in existing pheromone dispenser systems. Therefore, yield, sustainability, and cost of biosynthesis are the main considerations. From our experiments, we consider that dCas9EV2.1 mediated transient agroinfiltration is currently the best method for plant-based metabolite production. This provides the highest yields and enables predictable and equal expression from genes within multigene constructs. Further, when coupled with the relatively short timeline for production and the ability to rapidly prototype and implement new construct designs, this provides great potential for biomanufacturing. Further, the gene regulatory systems demonstrated here and developed as modular genetic elements for facile reuse, are not limited to controlling pheromone biosynthesis, but are broadly useful to the design of constructs for plant metabolic engineering.

## Methods

### Assembly of expression constructs

All constructs were assembled using the GoldenBraid (GB) cloning system (Sarrion-Perdigones et al., 2013; Vazquez-Vilar et al., 2017). Standardized DNA parts (promoters, coding sequences, and terminators) were cloned as Level 0 parts using the GoldenBraid (GB) domestication strategy described by Sarrion-Perdigones et al. (2013). Transcriptional units (Level 1) were then assembled in parallel, one-step restriction-ligation reactions and transformed into bacteria as previously described (Cai et al., 2020). Hierarchical stepwise assembly of transcriptional units into multigene constructs was achieved using binary assembly via BsaI or BsmBI-mediated restriction ligation as defined by the GB system. GB constructs employed in this study are provided in Supplementary Table S1 and have been deposited at Addgene.

### Transient expression in *N. benthamiana*

*N. benthamiana* plants were grown in a controlled environment room with 16 hr light, 8 hr hours dark, 22°C, 80% humidity, ~200 μmol/m^2^/s light intensity. Expression constructs were transformed into electrocompetent *Agrobacterium tumefaciens* GV3101. *A. tumefaciens* strains harboring the expression constructs were grown in LB medium supplemented with 50 μg/mL kanamycin or spectinomycin and 50 μg/mL rifampicin for 16 hours at 28°C/250 rpm. Overnight saturated cultures were centrifuged at 3,400 x g for 30 min at room temperature and cells were resuspended in infiltration medium (10 mM 2-(N-morpholino)ethanesulfonic acid (MES) pH 5.7, 10 mM MgCl_2_, 200 μM 3’,5’-Dimethoxy-4’-hydroxyacetophenone (acetosyringone)) and incubated at room temperature for 2-3 hours with slow shaking. Healthy plants (29-37 days old) with 3-4 fully expanded true leaves were infiltrated on the abaxial side of the leaf using a 1 mL needleless syringe and grown for five days in a growth chamber with 16 hr light, 8 hr hours dark at 22°C and 120-180 μmol/m^2^/s light intensity. Infiltrated leaves were treated with 2.5 mM copper sulfate by spray at three days post infiltration. The spray was applied to both the adaxial and abaxial surfaces of the leaf. All chemical compounds were purchased from Sigma-Aldrich (St. Louis, MO).

### Production of transgenic *N. benthamiana*

Constructs were transformed into *A. tumefaciens* strain LBA4404. Cells were collected from saturated cultures grown from a single colony and grown overnight to OD600 of 0.2 in TY medium (10gL^-1^ tryptone, 5gL^-1^ yeast extract, and 10gL^-1^ NaCl, pH 5.6) supplemented with 2mM MgSO4·7H2O, 200 μM acetosyringone and the appropriate antibiotics (Horsch et al., 1985). Leaves were harvested from immature, nonflowering plants, and surface sterilized. Leaf discs were cut using a 0.8-1.2cm cork borer and transferred to co-cultivation media (MS medium supplemented with vitamins enriched with 1mgL^-1^ 6-benzylaminopurine and 0.1mgL^-1^ naphthalene acetic acid). After 24 hours, the discs were incubated within the *A. tumefaciens* culture for 15 minutes and placed abaxial side down back on co-cultivation media. After two days, explants were transferred to selection medium (MS pH 5.8 supplemented with Gamborg’s B5 vitamins, 1mgL^-1^ 6-benzylaminopurine, 0.1mgL^-1^ naphthalene acetic acid, and 100 mgL^-1^ kanamycin). Explants were sub-cultured at 14-day intervals and shoots were transferred to rooting media (MS salts supplemented with Gamborg’s B5 vitamins and 100 mgL^-1^ kanamycin). Plantlets were transferred to soil and grown in a greenhouse (16 hlight, 24°C:8h dark, 20 °C).

### Quantification of reporter gene expression

Luciferase expression was detected using the Nano-Glo® Dual Luciferase® reporter assay system (Promega, Madison, WI, USA). Two 8 mm-diameter discs per infiltrated leaf were homogenized in180 μL passive lysis buffer (Promega) containing protease inhibitor (P9599, Sigma-Aldrich, Dorset, UK). Following incubation on ice for 15 min and centrifugation (100 × g, 2 min, 4°C), the supernatant was diluted to a 1:5 dilution. 10 μL of the dilution was mixed with 20 μL of passive buffer which was then mixed with 30 μL ONE-Glo™ EX Luciferase Assay Reagent (Promega) and incubated at room temperature for 10 min. LucF luminescence was detected using either a GloMax 96 Microplate Luminometer (Promega) or a Clariostar microplate reader (BMG Labtech, Aylesbury, UK) with a 10 s read time and 1 s settling time. LucN luminescence was detected from the same sample by adding 30 μL NanoDLR™ Stop & Glo® Reagent (Promega). After incubation for 10 min at room temperature, luminescence was detected as above. To calculate the proportion of expression from each reporter, luminescence from firefly luciferase (LucF) was scaled to the nanoluciferase (LucN) signal by an experimentally determined factor obtained from expression from single gene LucN and LucF constructs. Normalized (relative) expression levels of synthetic promoters were obtained as previously described (Cai et al., 2021) and are reported as the ratio of luminescence from the test promoter (LucN) to the calibrator promoter (LucF), normalized to the luminescence of an experiment control LucN/LucF expressed from calibrator promoters.

### Metabolite extraction and quantification

Standards, extraction methods and analysis of pheromone compounds were as previously described (Mateos-Fernández et al., 2021). Briefly, synthetic samples of Z11-16OH were obtained as described by Zarbin et al. (2007) and purified by column chromatography using silica gel and a mixture of hexane: Et2O (9:1 to 8:2) as an eluent. Acetylation of Z11-16OH was carried out using acetic anhydride (1.2 eq) and trimethylamine (1.3eq) as a base in dichloromethane (DCM). For biological samples, 8 mm leaf disks were snap frozen in liquid nitrogen and ground to a fine powder. 50 mg of frozen powder was transferred to 10 mL headspace vials and stabilized with 1 mL 5M CaCl_2_ and 150 μL 500mM EDTA (pH=7.5). Tridecane was added to a final concentration of 10 ppb for use as an internal standard and vials were bath-sonicated for 5 minutes. For volatile extraction, vials were incubated at 80°C for 3 minutes with 500 rpm agitation, after which the volatile compounds were captured by exposing a 65μm polydimethylsiloxane/ divinylbenzene (PDMS/DVB) SPME fiber (Supelco, Bellefonte, PA) to the headspace of the vial for 20 minutes. Volatile compounds were analyzed using a 6890 N gas chromatograph (Agilent Technologies, Santa Clara, CA) with a DB5ms (60m, 0.25mm, 1μm) J&W GC capillary column (Agilent Technologies) with helium at a constant flow of 1.2mLmin^-1^. Fiber was desorbed for 1 minute in the injection port at 250°C and chromatography was performed with an initial temperature of 160°C for 2 min, 7°Cmin^-1^ ramp until 280°C, and a final hold at 280°C for 6 minutes. All pheromone values were divided by the tridecane value of each sample for normalization. Alternatively, if tridecane was not added, pheromone values were normalized using the total ion count (TIC) of the corresponding sample (Wu and Li, 2016). For estimation of the yields, a calibration curve was constructed for each pheromone from a set of 7 peak areas ranging from 0.005 to 20 ppm, normalized with the tridecane values.

## Data Availability

Plasmids and their complete sequences have been submitted to Addgene (See Supplementary Table S1).

## Conflict of interest statement

None declared

## Author contributions

KK, EMG, DO and NP conceptualized the study. KK, EMG, RMF, SG and CT were responsible for the design and assembly of constructs. KK, EMG and CT conducted molecular and expression analyses of constructs. EMG, RMF, SG produced and analyzed transgenic lines. KK, EMG, RMF and SG conducted biochemical analyses of leaf extracts. All authors contributed to the analysis and visualization of data. DO and NP were responsible for supervision and funding acquisition. EMG, CT, and NP drafted the text. All authors contributed to revising and editing the text.

## Acknowledgements

We thank Yaomin Cai for advice and assistance with relative luminescence assays. We thank the John Innes Centre horticultural services team for assistance with plant growth and maintenance. We thank Ana Espinosa-Ruiz and Teresa Caballero-Vizcaino at the IBMCP’s metabolomic facility and Paul Brett at the John Innes Centre metabolomics facility for their help with GC-MS assays. We thank all members of the SUSPHIRE research consortium for helpful discussions throughout the project.

## Funding Statement

All authors gratefully acknowledge the European Research Area Cofund Action ‘ERACoBioTech’ for the support of SUSPHIRE (Sustainable Production of Pheromones for Insect Pest Control in Agriculture), which received funding from the Horizon 2020 research and innovation program under grant agreement No. 722361. KK, CT, and NP acknowledge the support of the UK Biotechnology and Biological Sciences Research Council (BBSRC) Core Strategic Program Grant to the Earlham Institute (Genomes to Food Security; BB/CSP1720/1) and grants BB/R021554/1 and BB/L014130/1. EMG, SG, RMF and DO acknowledge the support of grants PCI2018-092893 and PID2019-108203RB-100 from the Spanish Ministry of Economy and Competitiveness. EMG acknowledges a FPU grant (FPU18/02019) from the Spanish Ministry of Science, Innovation and Universities. RMF acknowledges a PhD grant (ACIF/2019/226) from the Generalitat Valenciana.

## Supplementary Data

*Supplementary Table S1*. Constructs used in this study

*Supplementary Figure S1*. Construct architecture influences expression from constitutive promoters.

*Supplementary Figure S2*. Copper inducible, CRISPR/Cas9-mediated control of pheromone biosynthesis.

*Supplementary Figure S3*. Transcription of dCas9:EDLL and MS2:VPR in T_1_ CBS:dCas transgenic plants.

*Supplementary Figure S4*. Pheromone biosynthesis in T_0_ transgenic *Nicotiana benthamiana* transgenics.

## Notes

### Competing Interest Statement

The authors have declared no competing interest.

### Summary of Updates

This version contains a revision to the data included in Figure 4. In addition new data are included (Figures 6 and 7). To integrate these, there are revisions to the text.

