## Supplementary Data for "Tunable control of insect pheromone biosynthesis in *Nicotiana benthamiana*"

**Supplementary Table S1** Constructs used in this study

| Plasmid Code | Contents | Addgene Code and Reference | Description |
|---|---|---|---|
| GB1203 | 35s:P19:nos | #68214 (Sarrion-Perdigones et al Plant Physiol. 2013 Jul;162(3):1618-31) | Transcriptional unit for constitutive expression of the silencing suppressor P19 driven by the 35s promoter. |
| 253 (pEPKKα1KN0253) | AtuNos+TMVΩ:CUP2:GAl4:35s | #187560 (this study) | Transcriptional unit for constitutive expression of CUP2:GAL4 driven by the nos promoter. |
| 021 (pEPCTΩSP0021) | CBS:LucF (forward):35s + CBS:LucN (forward) | #187561 (this study) | Module for copper-inducible expression of firefly luciferase and nanoluc luciferase driven by minimal synthetic promoters with binding sites for CUP2. |
| 022 (pEPCTΩSP0022) | CBS:LucN (forward):35s + CBS:LucF:35s (forward) | #187562 (this study) | Module for copper-inducible expression of nanoluc luciferase and firefly luciferase driven by minimal synthetic promoters with binding sites for CUP2. |
| 009 (pEPCTαKN009) | CBS:LucF:35s | #187563 (this study) | Transcriptional unit for copper-inducible expression of firefly luciferase driven by a minimal synthetic promoter with binding sites for CUP2. |
| 013 (pEPCTαKN013) | CBS:LucN (forward):35s | #187564 (this study) | Transcriptional unit for copper-inducible expression of nanoluc luciferase driven by a minimal synthetic promoter with binding sites for CUP2. |
| 017 (pEPCTΩSP0017) | 35s:LucF (forward):35s + 35s:LucN (forward) | #187565 (this study) | Module for constitutive expression of firefly luciferase and nanoluc luciferase driven by 35s promoters. |
| 018 (pEPCTΩSP0018) | 35s:LucN (forward):35s + 35s:LucF (forward) | #187566 (this study) | Module for constitutive expression of nanoluc luciferase and firefly luciferase driven by 35s promoters. |
| 019 (pEPCTΩSP0019) | 35s:LucF (forward):35s + 35s:LucN (reverse) | #187567 (this study) | Module for constitutive expression of firefly luciferase and nanoluc |

| | | | |
|---|---|---|---|
| | | | luciferase driven by 35s promoters. |
| 001 (pEPCTαKN001) | 35S:LucF:35S | #187568 (this study) | Transcriptional unit for constitutive expression of firefly luciferase driven by the 35s promoter. |
| 005 (pEPCTαKN005) | 35S:LucN:35S | #187569 (this study) | Transcriptional unit for constitutive expression of nanoluc luciferase driven by the 35s promoter. |
| GB UA 114 A | 35S:Gal4:PhiC31:35S | #187570 (Vazquez-Vilar et al Nucleic acids research vol. 45,4 (2017): 2196-2209) | Transcriptional unit for constitutive expression of Gal4:**Φ**C31 driven by the 35s promoter. |
| pEPKKα2KN0100 | 2xOpattB-min 35S:LucN:g7 | #154621 (Cai et al Nucleic Acids Res. 2020 Dec 2;48(21): 11845-11856) | Transcriptional unit for Gal4:**Φ**C31-activated expression of nanoluciferase. |
| pEPKKα2KN0101 | 4xOpattBt-min 35S:LucN:g7 | #154622 (Cai et al Nucleic Acids Res. 2020 Dec 2;48(21): 11845-11856) | Transcriptional unit for Gal4:**Φ**C31-activated expression of nanoluciferase. |
| pEPKKα2KN0102 | 6xOpattBt-min 35S:LucN:g7 | #154623 (Cai et al Nucleic Acids Res. 2020 Dec 2;48(21): 11845-11856) | Transcriptional unit for Gal4:**Φ**C31-activated expression of nanoluciferase. |
| pEPKKα1RKN0115 | AtuNos:TALE:35S | #187571 (Cai et al Nucleic Acids Res. 2020 Dec 2;48(21): 11845-11856) | Transcriptional unit for constitutive expression of a TALE driven by the nos promoter. |
| pEPKKα2KN0091 | 1xTALEbs-min 35S:LucNc:g7 | #154618 (Cai et al Nucleic Acids Res. 2020 Dec 2;48(21): 11845-11856) | Transcriptional unit for TALE-activated expression of nanoluciferase. |
| pEPKKα2KN0092 | 2xTALEbs-min 35S:LucNc:g7 | #154619 (Cai et al Nucleic Acids Res. 2020 Dec 2;48(21): 11845-11856) | Transcriptional unit for TALE-activated expression of nanoluciferase. |
| pEPKKα2KN0093 | 4xTALEbs-min 35S:LucNc:g7 | #154620 (Cai et al Nucleic Acids Res. 2020 Dec 2;48(21): 11845-11856) | Transcriptional unit for TALE-activated expression of nanoluciferase. |
| GB2085 | 35s:Ms2VPR:nos + 35s:dCas9:EDLL:nos | #160645 (Selma et al Plant Biotechnol J. 2019 Sep;17(9):1703) | Module for the expression of dCas9 fused to EDLL and Ms2 protein fused to VPR. |
| GB1724 | U626:gRNA4(pNOS) | #160621 (Selma et al Plant Biotechnol J. 2019 Sep;17(9):1703) | Transcriptional unit for a gRNA targeting the nos promoter with a MS2 recognition loop. |
| GB1838 | U6-26-1gRNA(pDFR) | #160625 (Selma et al Plant Biotechnol J. 2019 | Transcriptional unit for the expression of a g RNA targeting the DFR promoter with two copies of the MS2 |

| | | | |
|---|---|---|---|
| | | Sep;17(9):1703) | aptamer. |
| GB2513 | 35s:dCas9:EDLL:nos +35s:MS2:VPR:nos + U626:gRNA1 (pDFR) | #187803 (this study) | Module for constitutive expression of dCas9:EDLL, Ms2:VPR and a gRNA targeting the DFR promoter. |
| GB1024 | 35s:AtrΔ11:35s + 35s:HarFAR:35s | #187804 (this study) | Module for the constitutive expression of the Δ11 desaturase from *Amyelois transitella* and a fatty acid reductase from *Helicoverpa armigera*. |
| GB1022 | 35s:EaDAct:35s | #187805 (this study) | Transcriptional unit for expression of diacylglycerol acetyltransferase from *Euonymus alatus.* |
| GB3681 | 35s:ScATF1:35s | #187806 (this study) | Transcriptional unit for expression of alcohol O-acetyltransferase from *Saccharomyces cerevisiae* S288C, codon optimized for Nicotiana. |
| GB3682 | 35s:SpATF1-2:35s | #187807 (this study) | Transcriptional unit for expression of alcohol O-acetyltransferase from *Saccharomyces pastorianus* strain CBS 1483 chromosome SeVIII-SeXV, codon optimized for Nicotiana. |
| GB3683 | 35s:EfDAct:35s | #187808 (this study) | Transcriptional unit for expression of 1,2-diacyl-sn-glycerol:acetyl-CoA acetyltransferase from *Euonymus fortunei*, codon optimized for Nicotiana. |
| 678 (pEPKKΩ1SP0678) | 35s:TMVΩ:CUP2:GAl4: nos + CBSmin35s:AAtrΔ11:35 s + CBSmin35s: ATF1-2:mas + CBSmin35s:HarFAR:g7 | #187605 (this study) | Module for copper inducible expression of AtrD11, HarFAR and EaDAct. |
| GB3897 | minDFR:ATF1:mtb + minDFRHarFAR:pds + minDFR:AtrΔ11:dfr+ U626:gRNA1 (pDFR) | #187809 (this study) | Module for dCasEV2.1 activated expression of AtrD11, HarFAR and ScATF1 plus gRNA-1DFR. |
| GB4068 | nos:CUP2:GAL4:nos + CBS:dCas9:EDLL:nos + CBS: MS2:VPR;nos | #187810 (this study) | Module for the constitutive expression of Cup2:Gal4AD and copper-inducible expression of dCasEV2.1 (dCas9:EDLL and MS2:VPR). |
| GB4070 | nos:CUP2:GAl4:nos + U626:gRNA (DFR) + CBS:dCas9:EDLL:nos + CBS:MS2:VPR:nos | #187811 (this study) | Module for the constitutive expression of Cup2:Gal4AD and gRNA-1 DFR, and the copper-inducible expression of dCasEV2.1 (dCas9:EDLL and MS2:VPR). |
| GB3898 | minDFR:AtrΔ11:dfr + minDFR:HarFAR:pds + minDFR:ScATF1:mtb | #187812 (this study) | Module for dCasEV2.1 activated expression of AtrΔ11, HarFAR and ScATF1. |
| GB4356 | minDFR:AtrΔ11:dfr + minDFR:HarFAR:pds + minDFR:SpATF1-2:mtb | #187813 (this study) | Module for dCasEV2.1 activated expression of AtrΔ11, HarFAR and SpATF1-2. |
| GB4360 | minDFR:ScATF1:mtb + | #187814 | Module for dCasEV2.1 activated |

| | | | |
|---|---|---|---|
| | minDFR:AtrΔ11:dfr + minDFR:HarFAR | (this study) | expression of ATF1, AtrΔ11 and HarFAR. |
| GB4361 | minDFR:SpATF1-2:mtb + minDFR:AtrΔ11:dfr + minDFR:HarFAR:pds | #187815<br>(this study) | Module for dCasEV2.1 activated expression of ATF1-2, HarFAR and AtrΔ11. |
| GB4366 | minDFR:HarFAR:pds + minDFR:ScATF1:mtb + minDFR:AtrΔ11:dfr | #187816<br>(this study) | Module for dCasEV2.1 activated expression of HarFAR, ATF1 and AtrΔ11. |
| GB4367 | minDFR:HarFAR:pds + minDFR:SpATF1-2:mtb + minDFR:AtrΔ11:dfr | #187817<br>(this study) | Module for dCasEV2.1 activated expression of HarFAR, ATF1-2 and AtrΔ11. |

**Supplementary Figure S1.**

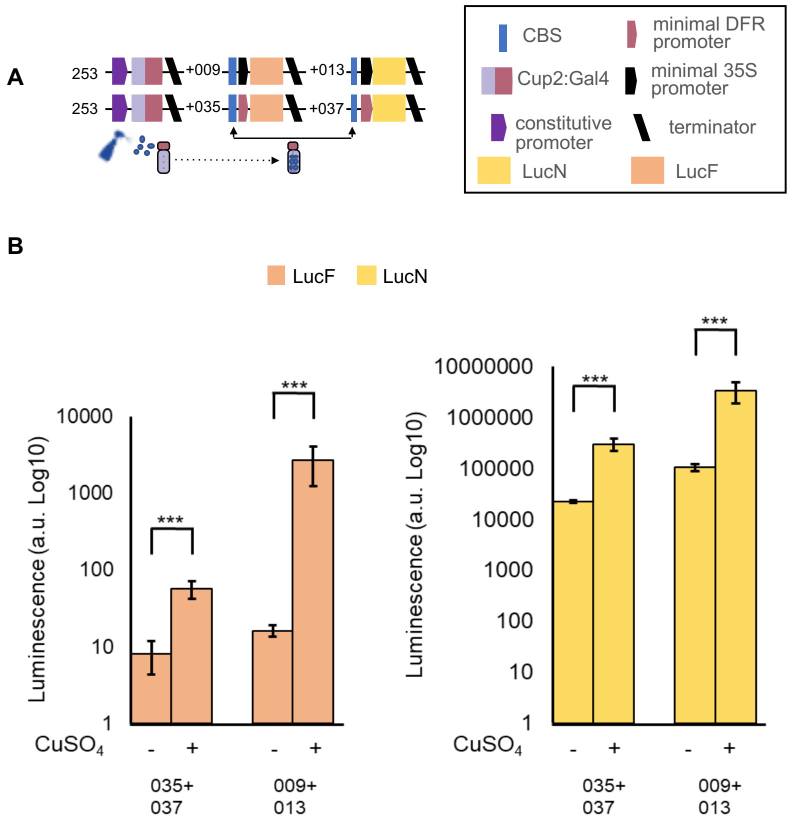


**Supplementary Figure S1.** Copper sulfate can induce Copper inducible promoters with minimal DFR or minimal 35S. (A) Schematics of plant expression constructs containing synthetic genes for copper inducible expression of firefly luciferase (LucF) and nanoluciferase (LucN). (B) Copper inducible promoters with minimal 35S or minimal DFR can be induced with copper sulfate (2.5 mM). Values shown are the mean and standard error of n=10 biological replicates (independent infiltrations) and differences were analyzed using pairwise Wilcoxon rank sum test with Benjamini-Hochberg correction (***P ≤ 0.001)

**Supplementary Figure S2.**

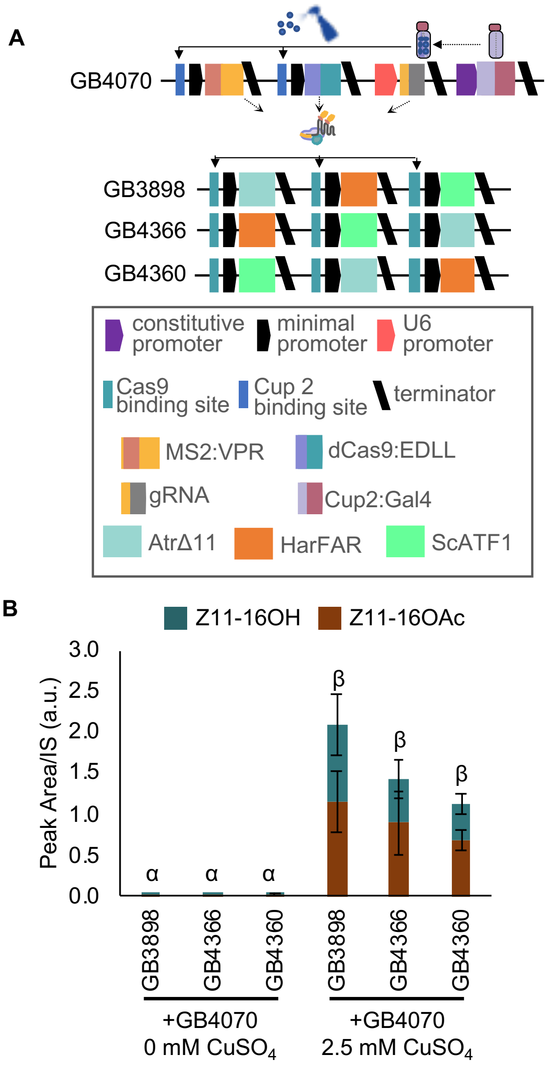


**Supplementary Figure S2. Copper inducible, CRISPR/Cas9-mediated control of pheromone biosynthesis.** (**A)** Schematic of plant expression constructs containing elements for copper inducible expression of the dCasEV2.1 transcriptional activator (above) and multigene constructs containing coding sequences for AtrΔ11, HarFAR and ScATF1, each assembled with a unique promoter with cognate target sequences for the gRNA (**B)** Application of $CuSO_4$ results in dCasEV2.1 mediated production of the pheromone components (Z11-16OH and Z11-16OAc). Values shown are the mean and standard error of n=3 biological replicates (independent infiltrations). Means followed by a common Greek letter (α, β) are not significantly different (one-way ANOVA with post-hoc Tukey HSD at the 5% level of significance).

**Supplementary Figure S3.**

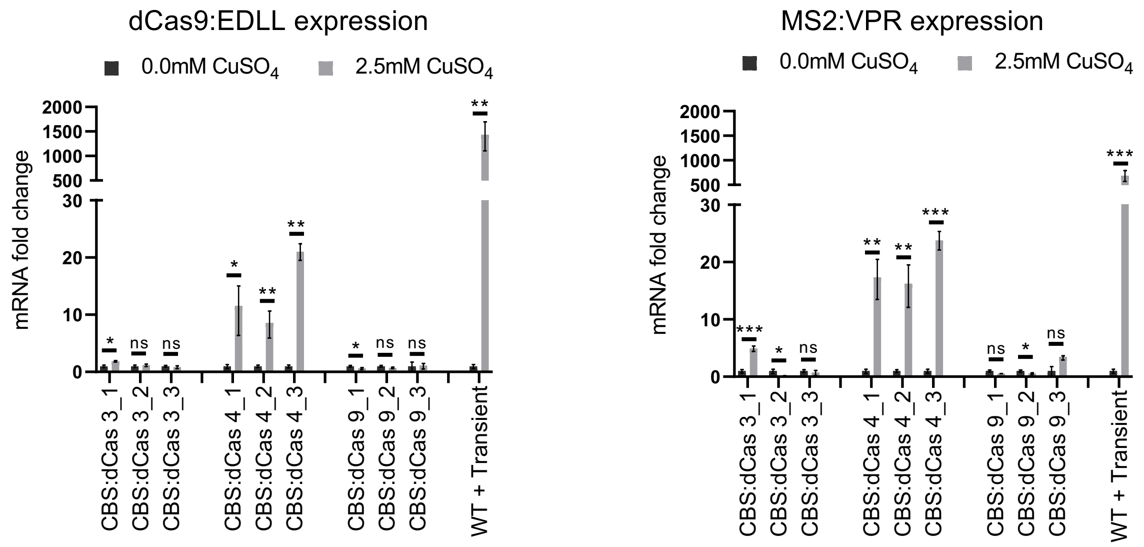


**Supplementary Figure S3.** Transcription of dCas9:EDLL and MS2:VPR in $T_1$ CBS:dCas transgenic plants. Three leaves from each of three transgenic progeny ($T_1$) of three independent transgenic T0 lines were selected and 0.0mM $CuSO_4$ or 2.5mM $CuSO_4$ applied each side of the midrib. Samples were collected from each plant 2 days after the induction (5dpi for the transient constructs). mRNA levels, relative to expression of the F-BOX gene (ΔCt) and and the ΔΔCt used to calculate fold change between copper concentrations. A WT plant infiltrated with the CBS:dCas module was included as a control. Error bars represents SD (n = 3). *P*-values were calculated using Student's t-test; **P* ≤ 0.05, ***P* ≤ 0.01, ****P* ≤ 0.001; ns= not significant.

### Supplementary Figure S4.

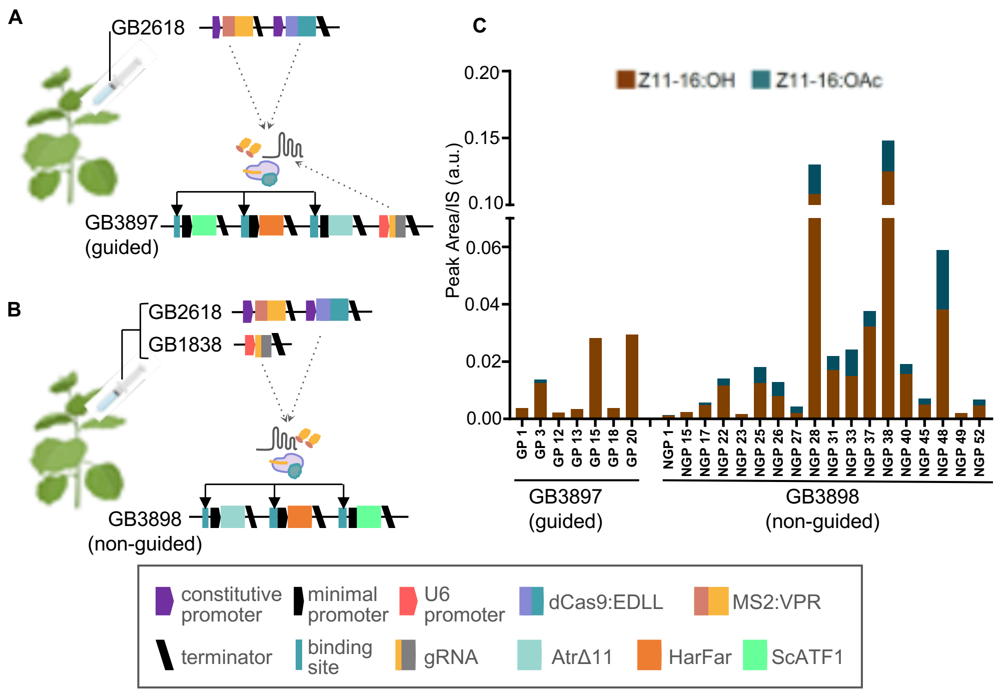


**Supplementary Figure S4. Pheromone biosynthesis in $T_0$ transgenic *Nicotiana benthamiana* transgenics**. Schematics of constructs used for expression of moth pheromones in transgenic lines containing the (**A**) guided pathway (sgRNA integrated) or (**B**) non-guided pathway (sgRNA infiltrated). (**C**) Pheromone levels obtained from $T_0$ transgenics infiltrated with constructs expressing regulatory elements. The figure includes images from Biorender (biorender.com).
